# Functional Ultrasound Imaging of the Human Spinal Cord

**DOI:** 10.1101/2022.08.06.503044

**Authors:** K. A. Agyeman, D.J. Lee, J. Russin, E.I. Kreydin, W. Choi, A. Abedi, V.R. Edgerton, C. Liu, V.N. Christopoulos

## Abstract

The integration of functional responses in the human spinal cord into the nervous system is not well understood. Herein we demonstrate the first in-human functional ultrasound imaging (fUSI) of spinal cord response to epidural electrical stimulation. fUSI is a minimally invasive neuroimaging technique that can record blood flow at a level of spatial and temporal precision not previously achieved in the human spinal cord. By leveraging fUSI and epidural electrical spinal cord stimulation in patients who underwent surgery, we recorded and characterized for the first-time hemodynamic responses of the human spinal cord to an electrical neuromodulatory intervention commonly used for treating pain, and increasingly used for sensory-motor and autonomic functions. We found that the hemodynamic response to epidural stimulation reflects a spatiotemporal modulation of the spinal cord circuitry not previously recognized. The impact of this analytical capability is significant for several reasons. It offers a mechanism to assess blood flow changes with a new level of precision which can be obtained in real time under *in vivo* conditions. Additionally, we demonstrate that fUSI can successfully decode the spinal cord state in a single trial, which is of fundamental importance for developing real-time closed-loop neuromodulation systems. Also, we show that spinal cord hemodynamic changes due to epidural electrical stimulation occur primarily at the level of small vessels. Overall, our work is a critical step towards developing a vital technique to study spinal cord function and understand the potential effects of clinical neuromodulation for spinal cord and other neurological disorders.

**One Sentence Summary:** The first in-human quantitative evaluation of spinal cord hemodynamics using functional ultrasound imaging (fUSI).

## Introduction

The spinal cord is a major sensorimotor integration center within the nervous system. It receives and monitors the kinetics, kinematics, and relative position of all parts of the body ^1–5^. Based on this constant directional flow of information the spinal networks provide a function in controlling most movements. Interruption of the bidirectional flow of information within the spinal cord secondary to traumatic injury or disease, at any level of severity, can lead to negative impacts ^6^. Deleterious effects include exaggerated reflex activity, chronic pain, partial to complete loss of motor and/or sensory function, bowel/bladder dysfunction, and adverse changes to sexual function ^7^. The cumulative effect of these dysfunctions severely impacts millions of individuals world-wide, leading to an immense burden on healthcare systems and national economies ^8–12^.

Despite the important role of the human spinal cord in sensory, motor, and autonomic functions, little is known about its functional architecture. While functional brain imaging (e.g., fMRI, stereoelectroencephalography [sEEG]) has been studied extensively (motor, sensory, language, memory), research on the functional anatomy of the spinal cord has been limited. The first fMRI studies of the human spinal cord were reported in the late 1990s ^13,14^. Since then, a growing but limited number of fMRI studies have attempted to reveal the functional organization of the spinal cord ^15–20^. Most importantly, the validity of BOLD signals as a hemodynamic proxy of spinal neural activity was only recently confirmed in non-human primates (NHPs), as signal variations were shown to correlate with electrophysiological activity (i.e., local field potentials) ^21^. The paucity of research could be attributed to technical challenges that arise when imaging the human spinal cord, including the small cross-sectional dimension of the cord (approximately 12 mm in diameter) ^22^, and susceptibility artifacts due to local magnetic field inhomogeneities created by interfaces between the surrounding bone, ligaments, soft tissues, and cerebrospinal fluid (CSF). Motion artifacts can also arise from the proximity to the thorax, lungs, and neck muscles ^23–25^. Optical and electrophysiological techniques that could potentially overcome some of these challenges face inherent limitations in scaling and penetration depth ^26,27^.

Given this context, the development of an imaging technology that can illuminate the functional architecture and hemodynamic responses of the spinal cord with high spatiotemporal resolution, sensitivity, and penetration depth is imperative. Until recently, no modality has been developed to successfully monitor the spinal cord to these criteria. Recently, functional ultrasound imaging (fUSI) was introduced as a breakthrough modality for large-scale recordings of neural activity ^28–31^. It provides a unique combination of great spatial coverage (∼ 10 cm), high spatiotemporal resolution (∼100 μm and up to 10 ms) and sufficient sensitivity (∼ 1 mm/s velocity) to detect hemodynamic changes of only 2% without averaging over multiple trials. While fUSI is a hemodynamic technique, its superior spatiotemporal performance and sensitivity offer substantially closer connection to the underlying neuronal signal than achievable with other hemodynamic methods such as fMRI. The first in vivo proof of concept for fUSI was established in 2011 by imaging cerebral blood volume (CBV) changes in the micro-vascularization of a rat brain during whisker stimulation ^29^. Since then, fUSI has been utilized to measure brain activity in freely moving rodents ^32,33^, awake and behaving NHPs ^34,35^, adult ^36,37^ and pediatric ^38,39^ human patients. By combining electrophysiology with fUSI in awake mice, a recent study showed that the power Doppler (pD) signal measured at frequencies below 0.3 Hz is strongly correlated with neuronal activity ^40^. This study established for the first time the direct association between neuronal activity and functional ultrasound signal. Recently fUSI was expanded to image rodent and swine spinal cord responses to different electrical response patterns ^41–44^, providing evidence that fUSI can detect C-fiber-evoked spinal cord hemodynamic responses elicited by activation of either natural noxious mechanical stimulations or electrically activated C fibers ^43^.

In this study we take a major leap in fUSI to quantify, for the first time, functional changes of the human spinal cord in response to epidural electrical spinal cord stimulation (ESCS). We monitored the hemodynamic activity of the spinal cord in 4 patients who underwent standard-of-care implantation of an ESCS device under general anesthesia for chronic back pain treatment. By acquiring fUSI of spinal cord activity at the level of the 10^th^ thoracic vertebra (T10) during ESCS (T8-9), we show that stimulation causes strong local hemodynamic changes in the spinal cord. This finding provides the first evidence that fUSI can capture and characterize evoked changes in spinal cord blood volume dynamics in humans.

We extend these results to accurately predict the effectiveness of a stimulation protocol at the single-trial level. By utilizing machine learning techniques to extract the most informative signals from the spinal cord that are associated with stimulation, we successfully predict the effects of electrical stimulation on spinal cord hemodynamics. Overall, our findings open new avenues to understand spinal cord function and provide a window into understanding the effects and potential of neuromodulation for neurological disorders.

## Results

To assess how electrical stimulation evokes specific changes in the spinal cord hemodynamics, we acquired fUSI images from 4 patients who underwent ESCS surgery (T10 partial laminectomy) for chronic back pain treatment (Fig. 1A-C). We used a miniaturized 15-MHz linear array transducer, inserted through a laminar opening onto the dura of the spinal cord with transverse field of view (Fig. 2A). The recordings produced pD functional ultrasound images of the spinal cord with a spatial resolution of 100 μm x 100 μm in-plane, slice thickness of about 400 μm and field of view (FOV) 12.8 mm x 10 mm. The power Doppler-based functional ultrasound image from a typical patient (*patient 1*) (Fig. 2B) illustrates for the first time that fUSI can capture the anatomical vascularization of the human spinal cord.

**Figure 1.**
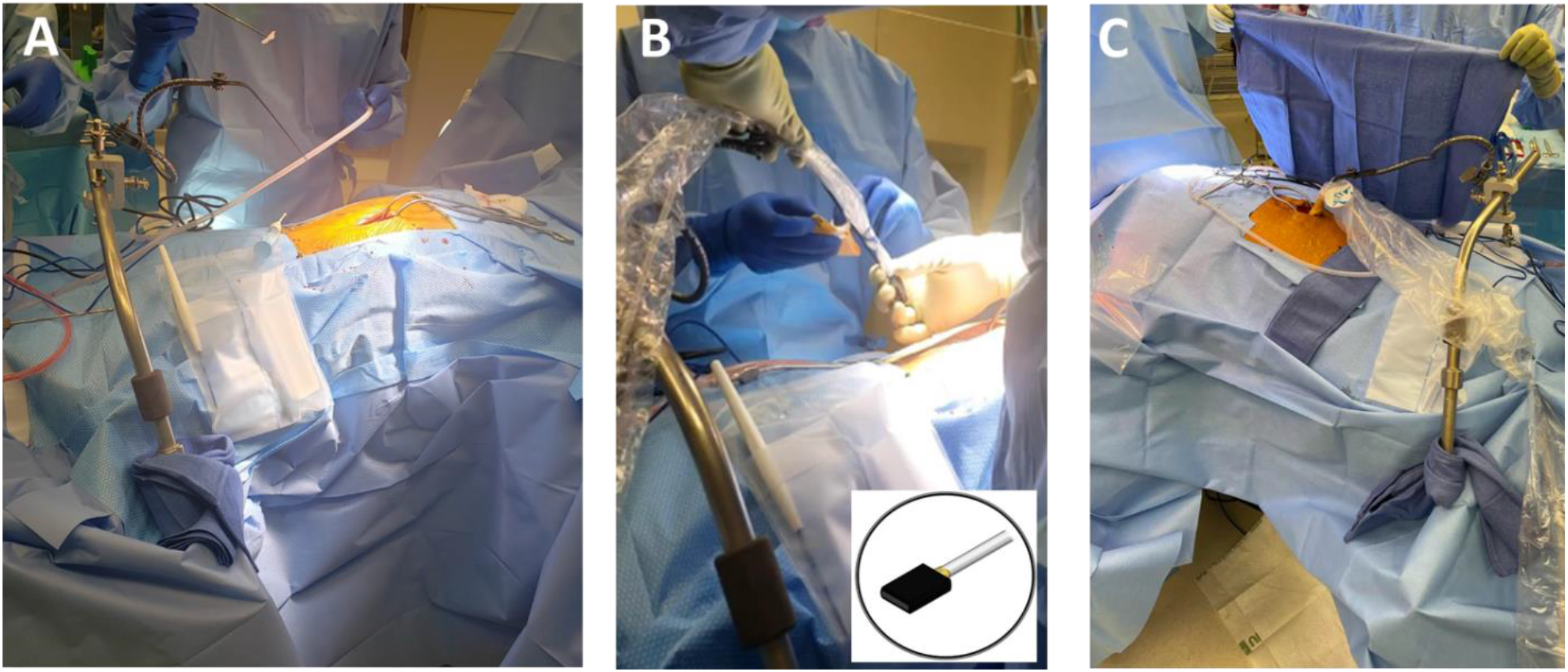
Patients and surgical procedure. **A)** Patients undergo a T10 partial laminectomy for standard-of-care implantation of an epidural spinal cord stimulator ESCS (PentaTM model 3228), under general anesthesia. **B**) A 15 MHz functional ultrasound probe (inner panel) is inserted into a sterilized cover to reduce the level of microbial contamination. Probe is mounted onto a retractor surgical arm. **C)** fUSI images being acquired on a transverse plane through the laminar opening with transducer placed onto the spinal cord dura.

**Figure 2.**
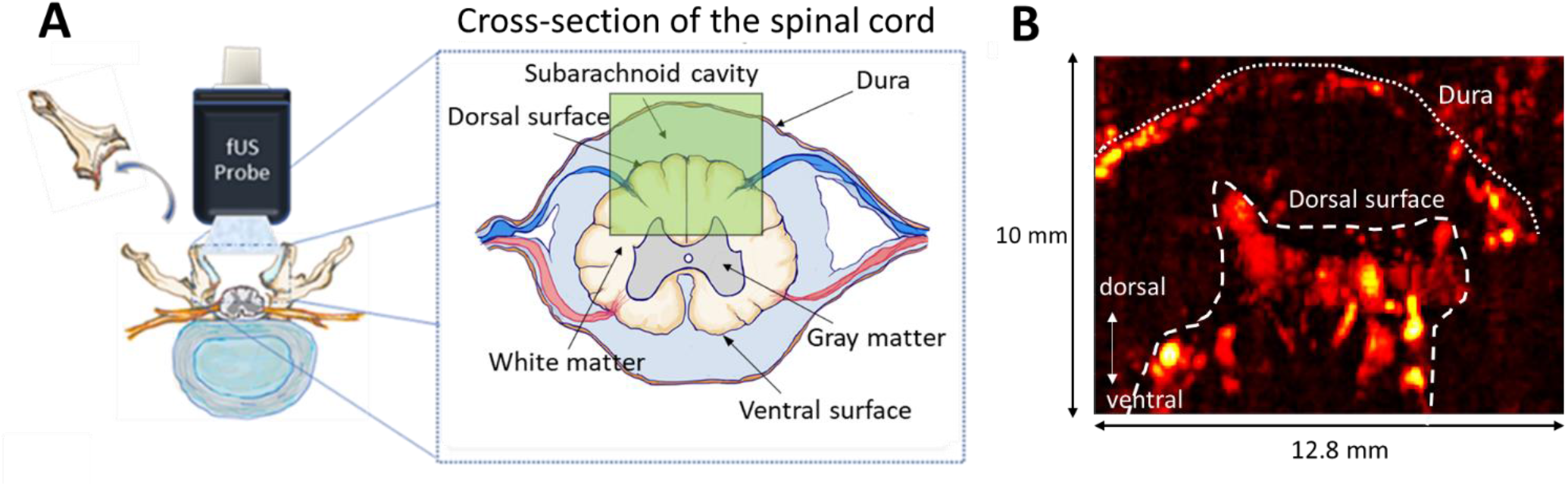
Functional ultrasound imaging of the spinal cord in a transverse plane. **A)** Schematic representation of spinal cord fUSI through a laminar window and cross section of spinal cord anatomy. The green area illustrates approximately the field of view of fUSI acquisition *(patient 1)*. **B**) Power Doppler-based functional ultrasound image showing the transverse section of the spinal cord of a typical patient *(patient 1)*. The field of view includes the dura and the dorsal surface of the spinal cord (white discontinuous lines).

### Hemodynamic responses induced by ESCS

To resolve ESCS-specific hemodynamic changes, we employed a stimulation protocol in which electrodes were placed to span the T8-9 spinal interspace location to deliver 10 ON-OFF cycles, with each cycle containing a 30 s ON period (stimulation ON) and a 30 s OFF period (stimulation OFF). Burst frequency of 40 Hz with 300-ms pulse width and 6 mA amplitude were utilized. The same ESCS protocol was used for all patients. A typical hemodynamic response to ESCS in a single stimulation cycle (30 s OFF, 30 s ON) of patient 1 is illustrated in Fig. 3 (see movie S1 in the supplementary materials). The mean fUSI background grayscale pD image is overlaid with color coded measured changes of the spinal cord blood volume (ΔSCBV) from the baseline activity (i.e., average fUSI activity for 6 seconds preceding the stimulation onset). ESCS causes regional changes in blood flow – some regions showed an increase (reddish) of blood flow whereas other regions exhibited a decrease (blueish) of blood flow after turning on the stimulator (Fig. 3). Note that once the stimulator is turned off, blood flow response returns progressively to the baseline condition. These localized dynamics in blood flow may reflect continuous changes in the physiological states among networks during ESCS. To the best of our knowledge, this is the first study that depicts hemodynamic responses of the human spinal cord to ESCS.

**Figure 3.**
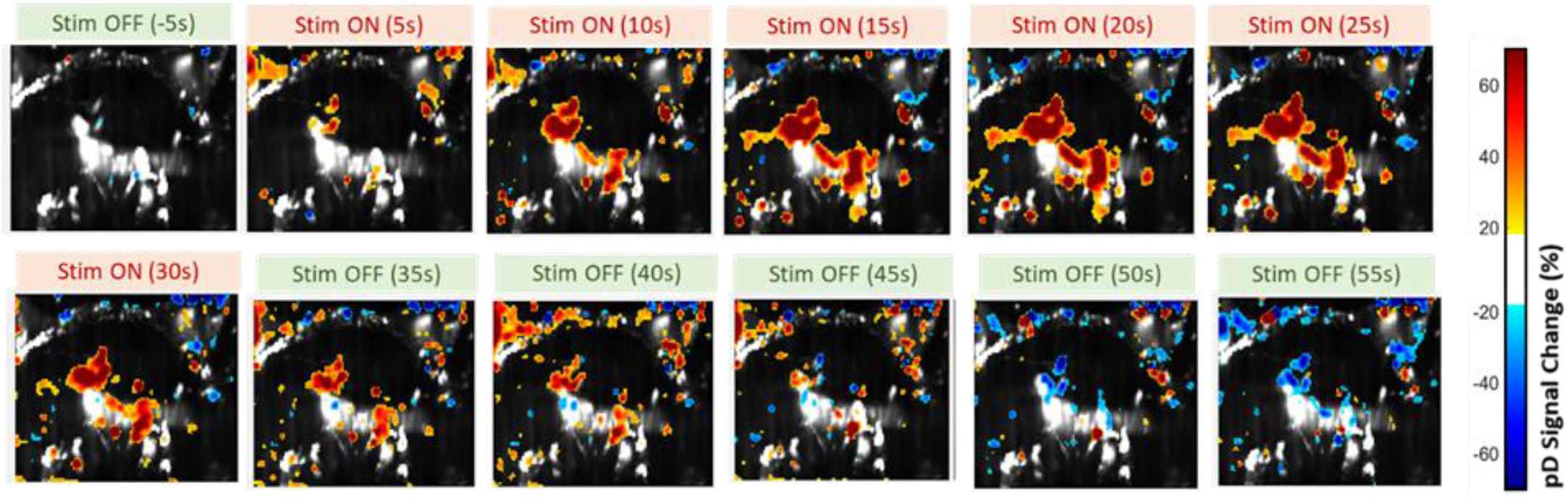
Hemodynamic response of the spinal cord induced by epidural electrical spinal cord stimulation (ESCS) in a typical patient. Spatiotemporal spinal cord blood volume change (ΔSCBV) greater than 20% relative to baseline, color coded on the power Doppler (pD) images at several time points starting 5 seconds before onset of the stimulation *(patient 1)*. Baseline activity was computed as the average fUSI signal for 6 s prior to stimulation onset.

To better understand how ESCS evokes specific changes in spinal cord hemodynamics, we compute statistical parametric maps (SPM) based on one-sided Student’s t test with false discovery rate correction (FDR). The SPMs identify regions recorded within the spinal cord that are strongly affected by electrical stimulation. They provide spatial visualization of spinal cord regions where the SCBV during stimulation ON is significantly different compared to the SCBV during stimulation OFF (supplementary Figure S2). It could be argued that the effect of the electrical stimulation to the hemodynamic signal is carried over to the stimulation OFF trials. There is evidence that electrical stimulation can result in prolonged clinical and physiological changes after the stimulation is turned off ^45,46^; however, the exact length of these changes in spinal cord stimulation is not well understood. Thus, analyzing the entire OFF period, SPMs may not accurately visualize spinal cord areas which are strongly affected by stimulation, since the hemodynamic signal can take up to 20 s to return to baseline activity after ceasing the stimulation (i.e., stimulation OFF trials) (see following analysis in Figs. 4B and 4C). To minimize the potential washout effect of the electrical stimulation, we computed the SPMs using only the last 6 s of the stimulation OFF period (Fig. 4A). The results showed that the spatial patterns of SCBV changes are preserved in all patients even if we do not correct for washout effects from the electrical stimulation. In other words, SPMs are similar regardless of using 30s (total period) or the last 6 s of the stimulation OFF trials.

**Figure 4.**
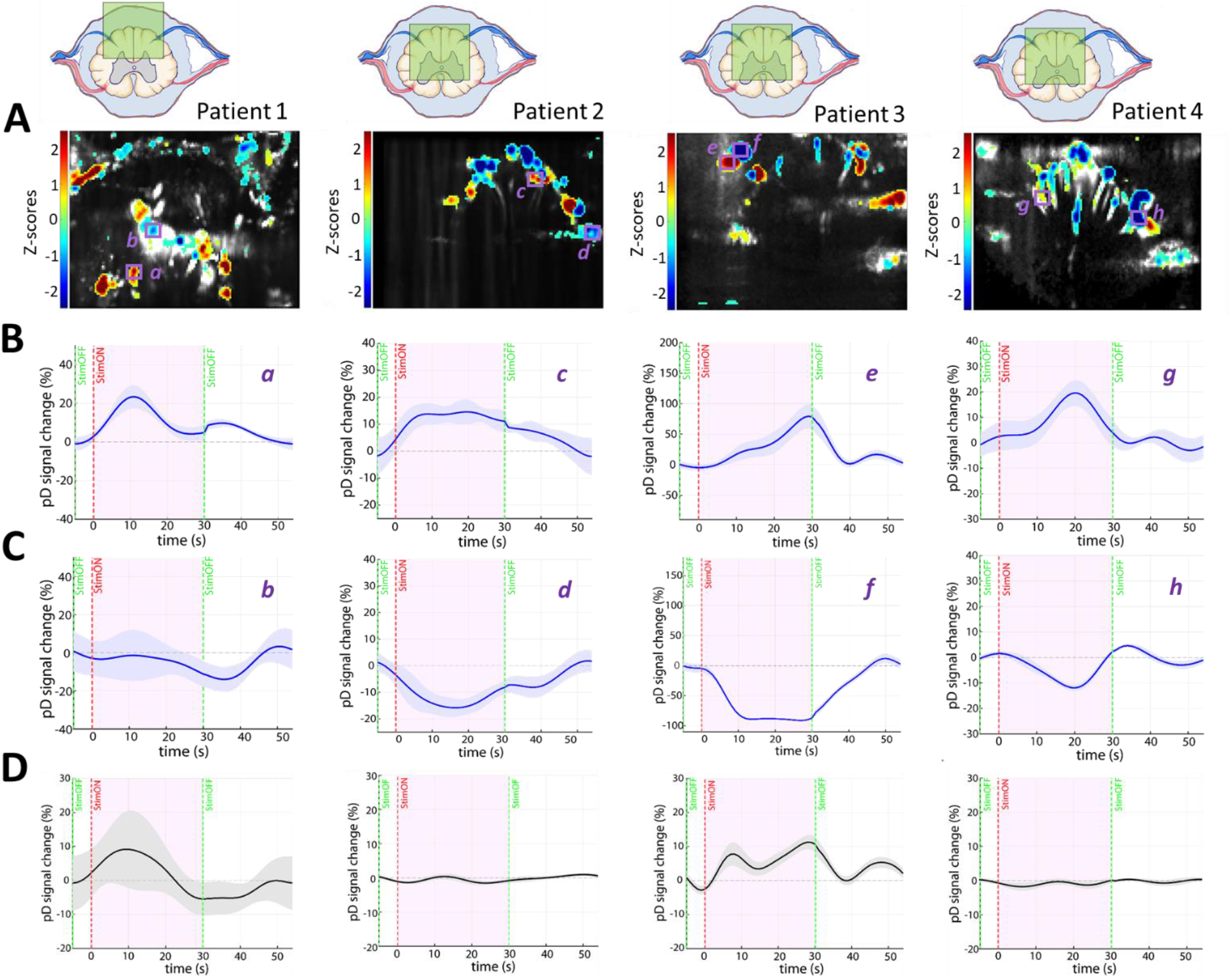
Statistical parametric map (SPM) and event-related average (ERA) waveforms *(Four patients)*. **A**) Statistical map shows localized areas with significantly higher SCBV with respect to baseline (one-sided t test of area under the curve, p < 0.05, false discovery rate [FDR] corrected for number of pixels in image). The top panels illustrate approximately the field of view of the fUSI acquisition for each patient. **B**) ERA waveforms of selected regions of interest (ROIs) [as indicated with boxes and letters in SPMs – panels **A**] that exhibit strong increased changes of blood perfusion after turning on the stimulator. Notice that some ROIs exhibit stronger changes of blood perfusion than others. **C**) ERA waveforms of selected regions of interest (ROIs) [as indicated with boxes and letters in SPMs – panels **A**] that exhibit strong decrease change of blood perfusion after turning on the stimulator. The background grayscale vascular images illustrate the average power Doppler signal of the spinal cord across the acquisition. **D**) ERA waveforms of the *global* SCBV changes of the 4 patients. The *global* SCBV is determined as percentage change in power Doppler (pD) from baseline of the total recorded spinal cord region recorded.

Additionally, we generated event related average (ERA) waveforms of the ΔSCBV as a percent change from the baseline activity to elucidate the temporal patterns of the hemodynamic changes of the spinal cord during ESCS. The results show that ESCS causes strong regional changes in blood flow in all 4 patients (Figs. 4B & 4C). Notably, we observed rising (Fig. 4B) and falling (Fig. 4C) ERA curves with regional-dependent peak responses and peak time induced by ESCS for all patients. This is followed by a subsequent return to baseline before or immediately after stimulation is ceased. Note that fUSI is currently limited to acquisition in a 2D imaging plane, since ultrafast Doppler relies on a limited number of channels that drive 1D ultrasonic linear probes. Therefore, it is challenging to acquire the same fUSI plane across experimental sessions and patients. However, although the imaging planes vary across patients – specifically, the depth of imaging plane in patients 2, 3 and 4 are deeper (approximately 6 mm) than the depth of the imaging plane in patient 1 (Fig. 4A top row) – the spatial and the temporal patterns of the SCBV changes induced by ESCS remain consistent across patients. This observation highlights the strength and robustness of fUSI to overcome the potential to image different 2D slices between experimental sessions and patients. Together these results provide the first evidence that fUSI can capture and characterize spinal cord hemodynamic changes induced by ESCS in humans.

### Single-trial decoding analysis

The capability to evaluate the effects of ESCS on spinal cord hemodynamics without having to perform multiple stimulation trials, and modify the protocol if needed, could have tremendous translational clinical implications. Intuitively, the simplest way to evaluate whether a stimulation protocol evokes hemodynamic changes of the spinal cord would be to measure SCBV changes (i.e., %pD signal change from the baseline) of the whole recorded spinal cord region (named global ΔSCBV). However, this seems to be problematic since the hemodynamic changes evoked by ESCS vary across spinal cord regions. Blood flow in certain regions takes longer to increase from baseline, whereas other regions exhibit faster induced blood flow changes. Additionally, while some regions show increases, others show decreases in blood flow during ESCS. The ERA waveforms of the global ΔSCBV of the 4 patients (Fig. 4D), is consistent with our prediction that electrical stimulation does not always cause clear changes of the *global* SCBV, even averaging across multiple trials. This suggests that the global ΔSCBV may not reflect a reliable measurement to assess whether a stimulation protocol causes hemodynamic changes on the spinal cord. Hence, a more sophisticated analysis is needed to detect hemodynamic changes induced by electrical stimulation.

One of the great advantages of fUSI is the ability to detect regional hemodynamic changes of only 2% without averaging over multiple trials ^35^. Thus, we explore whether fUSI can predict the effect of ESCS on hemodynamics of the spinal cord within a single trial. To do so, we implemented a class-wise principal component analysis (cPCA) to extract effective discriminant features (Fig. 5). That is, we reduced dimensionality of the fUSI images by selecting spinal cord regions that exhibit the highest pD signal difference between stimulation ON and stimulation OFF trials and optimally discard shared information – i.e., noise between class 0: stimulation OFF and class 1: stimulation ON (Fig. 5). cPCA has been used to reduce sparsity and dimensionality while maintaining enough components to retain over 90% variance in the data (see Materials and Methods section for more details). It is ideally suited for discrimination problems with large dimension and small sample size including natural and biomedical images ^47,48^. Finally, to predict the epidural stimulation induced states of the spinal cord, we used a linear discriminant analysis (LDA), where 75% of the event aligned (stimulation ON and stimulation OFF) trials fUSI images were randomly selected to train the model with 25% hold-out test dataset. The results of bootstrapping (100 iterations) and cross-validated accuracy show that the performance accuracy of the decoding algorithm is 84.0 ± 11.4 % across the 4 patients (MEAN ± STD).

**Figure 5.**
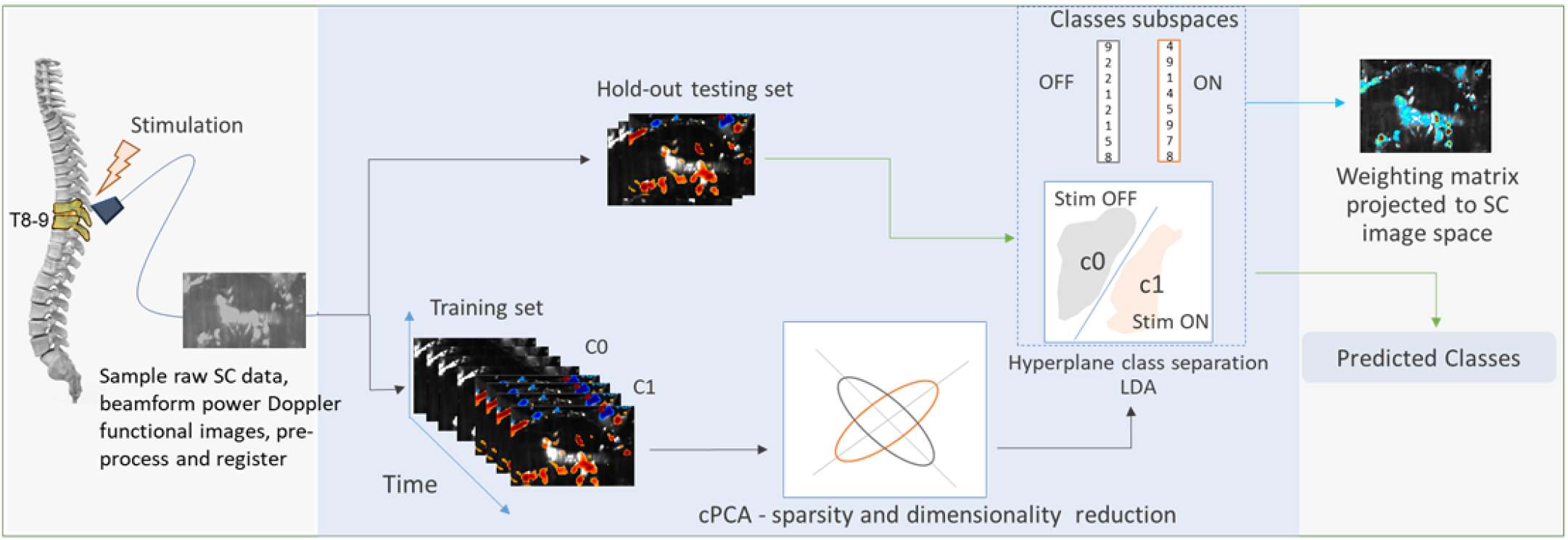
Flowchart for single-trial stimulation state decoding from the fUSI signal. Training images were separated from testing images based on the cross-validation technique used – i.e., 75% training data and 25% test data from each class. The state of the spinal cord was decoded from single trials based on a cPCA dimensionality reduction algorithm and a linear discriminant analysis classification model built by the training pD spinal cord imaging data with the corresponding class labels (i.e., 0: OFF stimulation, 1: ON stimulation).

To analyze the temporal evolution of the stimulation-related information in the spinal cord, we attempted to decode the state of the spinal cord across the stimulation time. We used a 1-s incremental window of data to train the classifier and then attempted to decode the state of the spinal cord using pD test images acquired within that window. For example, for t = 10 s, we trained the classifier using pD images of both classes from t=1-10 s (where t = 1 s corresponds to one second after stimulation onset for ON trials and after ceasing the stimulation for OFF trials), and then, we attempted to decode the state of the spinal cord using pD images acquired within that 10 s window. The resulting average cross-validation accuracy curve shows that the classification performance increases with the stimulation time (i.e., training window), reaching a plateau at around 15 s after stimulation onset, but then decreases when more than 25 s imaging data is accumulated to train the classifier (Fig. 6A). These results suggest that the stimulation effects to the hemodynamic signal of the spinal cord start decreasing before turning off the stimulator, and therefore the functional images acquired after that time and used to train the classifier may reduce the decoding accuracy of the spinal cord state. Distinct spatial locations within the spinal cord encode stimulation-related information as reflected in the variable weighting assigned to each voxel in the decoding algorithm (Fig. 6D). Notice that the spinal cord regions that encode stimulation-related information vary with stimulation duration (i.e., the amount of data used to train the classifier). Together, these findings suggest that the amount of data used to train the classifier is critical for the decoding accuracy of the spinal cord state. As such, the next step is to determine the time point after the stimulation onset that the classifier achieves the maximum decoding accuracy for a given amount of training data (i.e., training window). This analysis aims to find the time point after stimulation onset in which the pD images provide the best discrimination between the two classes. To do so, we used a 1-s incremental window of data to train the classifier as before, but now we attempted to decode the state of the spinal cord using pD test images acquired at a specific time point *t* within the time window that the classifier was trained. For instance, when the classifier is trained using pD images within a window t=1-5s, the decoding accuracy is evaluated at time t=1, 2, 3, 4, and 5s. Fig. 6B depicts the average decoding accuracy across patients for pD images (i.e., test data) acquired at specific time instants within the training window. The results show that the best decoding accuracy (98.3%) is achieved at t = 17 s after turning on the stimulation for a classifier trained using t=1-24s pD images (training window). Overall, these findings provide strong evidence that fUSI can evaluate the effectiveness of a stimulation protocol within single trials, opening new avenues in closed-loop neuromodulation technologies.

**Figure 6.**
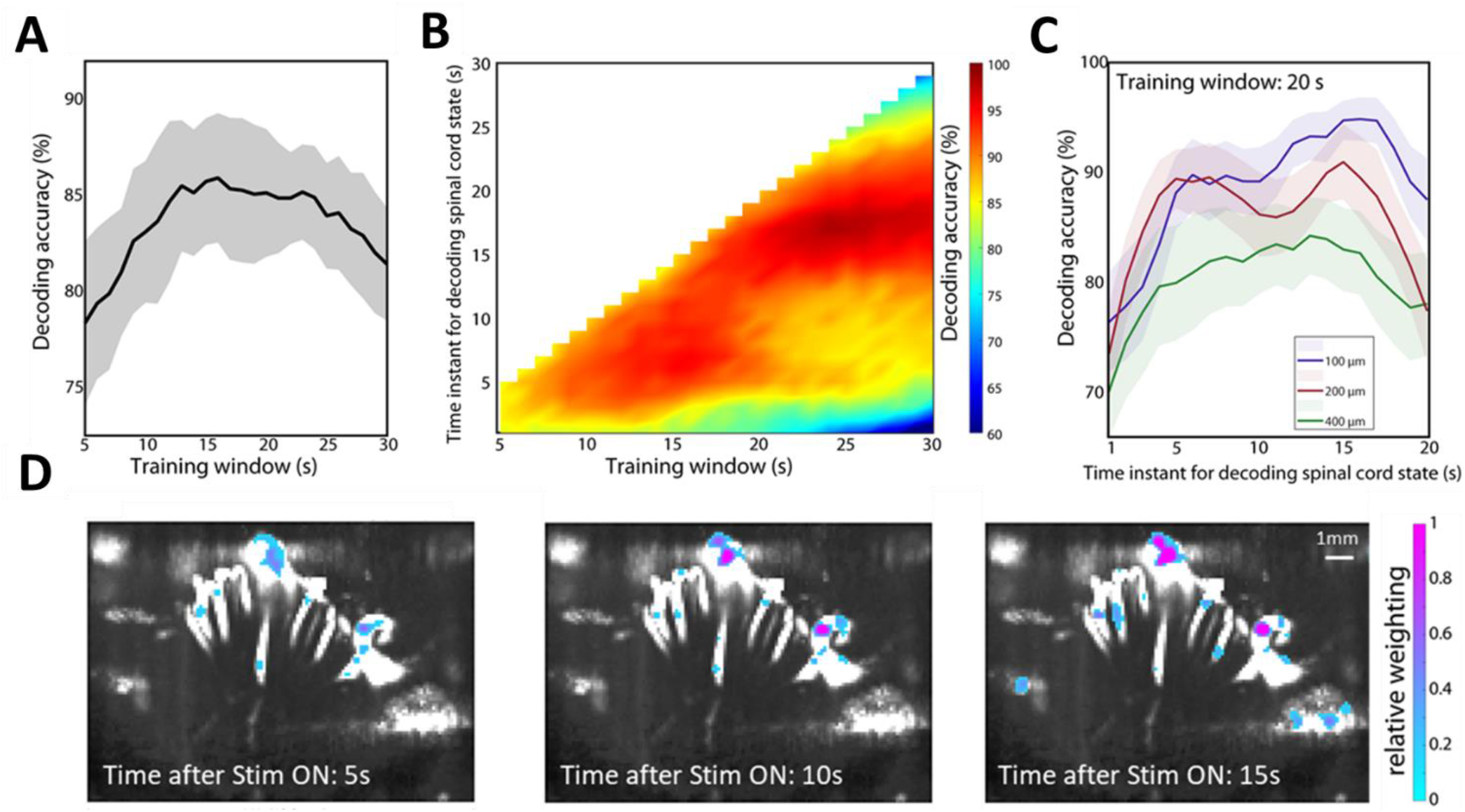
Single-trial decoding of the spinal cord state. **A)** Average decoding accuracy across patients as a function of the amount of data used to train the classifier – i.e., time after the stimulation onset - training window. **B**) Decoding accuracy of the spinal cord state using pD images acquired at specific time points after turning on the stimulation (with time step 1 s) as a function of the amount of data used to train the classifier (i.e., training window). For instance, we train the classifier using a set of pD images between t=1-5 s and evaluate the decoding accuracy of each second – 1s, 2s, 3s, 4s, and 5s. **C**) Decoding accuracy of the spinal cord state for pD training images acquired between 1s and 20s after turning on the stimulator – i.e., training window of 20s. Different traces correspond to different spatial resolutions (100 μm, 200 μm and 400 μm) of the pD images. **D**) Representative decoder weighting maps (patient 4). The top 10% most heavily weighted voxels are illustrated as a function of space and time after the stimulation onset, overlaid on the vascular map.

### Vascular signal and information content in the human spinal cord

Compared to other functional neuroimaging modalities, fUSI provides unprecedented resolution and sensitivity. To evaluate the benefits of enhanced spatial resolution in studying the human spinal cord, we attempted to decode the spinal cord state using 1-20 s window of pD images to train the classifier while decreasing the resolution of the images. We resized the images using a nearest neighbor interpolation approach to achieve spatial resolution of 200 μm and 400 μm – the resolution of the original pD image is 100 μm. We then used the downsized images, which contained fewer pixels than the original images, to decode the spinal cord state. The results show that the decoding accuracy decreases as voxel size increases (Fig. 6C) proving the importance of high spatial resolution in studying the human spinal cord.

We also hypothesized that the most informative power-Doppler content, useful for decoding the spinal cord state, is primarily located in small vessels. The hemodynamic response is believed to start in parenchymal arterioles and first-order capillaries ^49^. We tested this hypothesis by rank-ordering all voxels based on mean pD intensity and segmenting them into deciles. This resulted in spatial maps of ranked deciles (Fig. 7A), which indirectly correlate with spinal cord blood flow and vessel size – deciles 1-5 capture small vessels, while deciles greater than 6 correspond to large arteries. We subsequently utilized each decile to independently classify the spinal cord state (i.e., stimulation OFF vs. stimulation ON) and then evaluated for corresponding normalized classification accuracy. In this analysis the classifier was trained using pD images acquired during the total acquisition period – i.e., time window 1-30s. Peak accuracy (0.93 ± 0.018; MEAN ± SEM) was observed when the fourth mean pD ranked decile was used to classify the spinal cord state across patients (Fig. 7B) – a 10.24% increase in decoding accuracy, compared to when the entire spinal cord image is used. The accuracy declines when large-artery deciles (> 6) are used for the classification analysis. The results are consistent with our hypothesis that the most informative pD hemodynamic content is linked to smaller vessels (i.e., arterioles and capillaries). We are the first to show this outcome in the human spinal cord. This observation is a unique outcome that could generate important cues in optimizing human spinal cord neuromodulation, similar to observations in recent NHP ^50^, rodents ^51^, and ferrets ^52^ brain studies.

**Figure 7.**
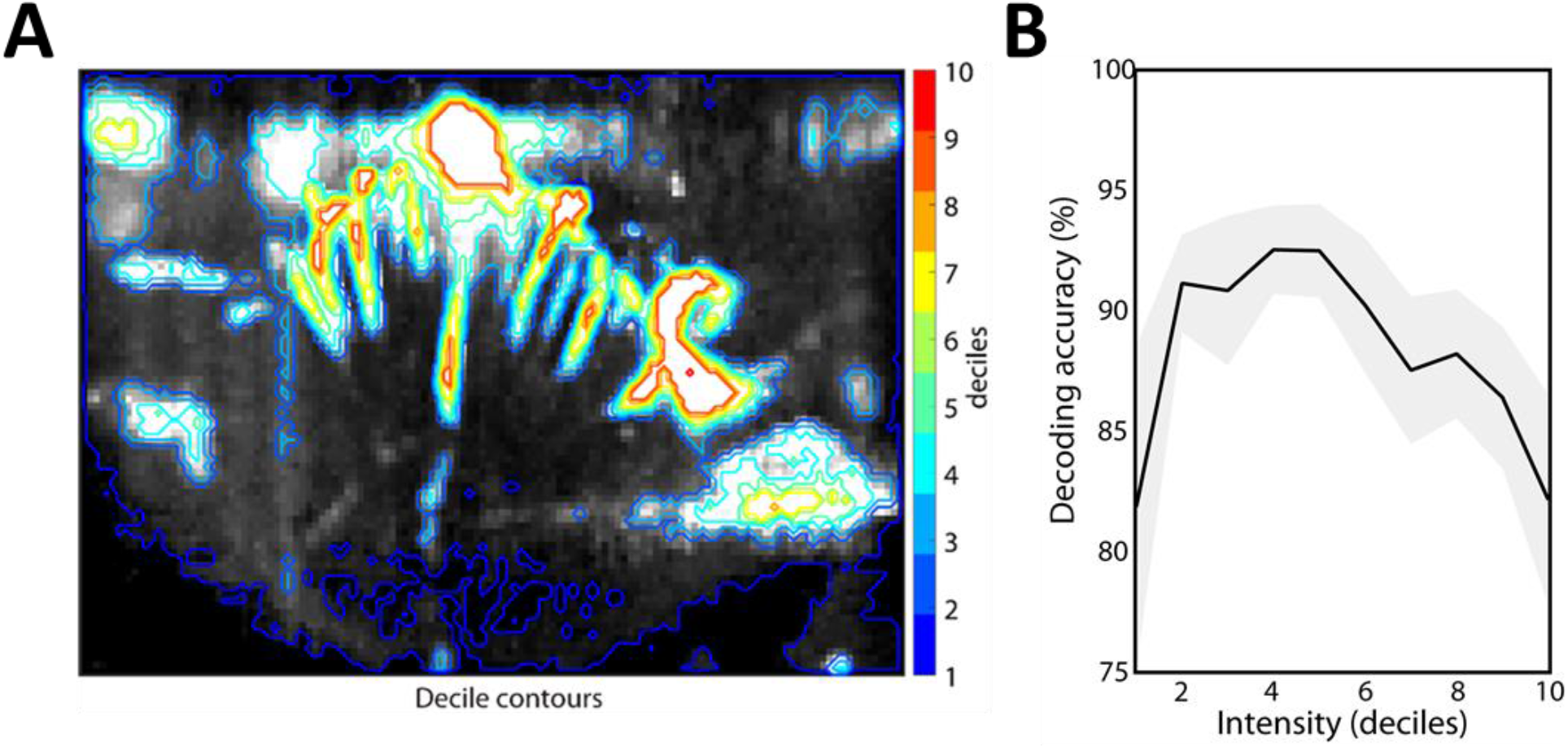
Effects of mean power Doppler in single-trial decoding analysis. **A)** A vascular map of patient 4 with contours dividing the image into deciles of mean power Doppler (pD) intensity. **B**) Decoding accuracy (Stimulation ON vs. Stimulation OFF) as a function of the pD intensity – i.e., deciles. Decoding accuracy peaks around quantile 4, which mostly contains small vasculature within the spinal cord. Smaller vessels and primary unit vasculature (i.e., deciles 1 and 10, respectively) carry less information about the effect of the ESCS to spinal cord blood flow. Black trace represents mean decoding accuracy and shaded area represents standard error (SEM) across all 4 patients.

## Discussion

### General

To date, functional imaging of the human spinal cord has been limited. Here, we utilize fUSI to image physiological manipulation of the spinal cord through ESCS. Much of the current knowledge on how the spinal cord processes sensorimotor information and generates actions comes from animal research due to the complexity and ethical issues related to access to the human spinal cord. In fact, traditional studies of human spinal cord functions have predominantly relied upon indirect electrophysiological measurements, through coupling of electromyogram recordings with electrical and/or magnetic stimulation of peripheral nerves, cerebellum, and cortical brain regions ^53–55^. While these studies have significantly improved our understanding of the sensorimotor processes in the spinal cord, they fall short of providing critical understanding of how sensory ensembles are translated into information about functional spinal networks. Recently, a handful of fMRI studies aimed to provide a large-scale view of the functional organization of human spinal circuits (for review, see ^56^). However, most of these studies focused only on the cervical level of the spinal cord ^14,19,57–61^ because imaging the thoracic and lumbar segments is more challenging due to a smaller cord diameter and greater physiological noise, such as breathing. To overcome the technical challenges associated with spinal cord fMRI, the current study uses functional ultrasound imaging (fUSI) to monitor the hemodynamics of the spinal cord at the thoracic level. We show for the first time that fUSI can capture and characterize hemodynamic changes in the spinal cord induced by electrical stimulation applied epidurally, rostral to the recording site but at the most rostral segments of the lumbosacral segments where the spinal locomotor networks are located, including interneurons that contribute to central pattern generators. These control much of the movement of the lower limbs, bladder, bowel and sexual functions. Our results demonstrate that electrical stimulation causes regional changes in spinal cord hemodynamics, with certain regions that exhibit significant increases and others significant decreases in blood flow. By taking advantage of the unprecedented spatiotemporal resolution and sensitivity of fUSI, we show that this modality can accurately predict the effects of a stimulation protocol within a single trial.

### Functional ultrasound imaging to guide neuromodulation treatment

Detecting and characterizing the effects of epidural spinal cord stimulation (ESCS) is crucial to unravelling the complex mechanisms of action subserving chronic pain suppression. Chronic back pain represents the leading cause of disability worldwide, affecting an estimated 540 million people at any given time. Traditional treatments such as surgery, opioid analgesics, and physical therapy ^62–66^ often prove ineffective, or in some cases, increase pain and/or worsen quality of life ^67,68^. Over the past couple of decades ESCS has emerged as a neuromodulation tool to treat chronic pain conditions that are refractory to traditional therapies ^69^. It involves surgical implantation of a small array of electrodes into the epidural space, akin to a pacemaker that deliver electrical pulses to the spinal cord for masking pain before reaching the brain ^70^. While spinal cord stimulation is effective for many patients, it is often associated with suboptimal efficacy and short-lived therapeutic effects ^71^. It is estimated that nearly 45% of patients fail to experience significant improvement in their pain after stimulator implantation. Some patients do not tolerate the persistent perception of paresthesia, while others do not obtain adequate pain relief from conventional ESCS. Predicting which patients will not have a resolution of pain remains challenging, mainly because the mechanism of action (MOA) by which ESCS alleviates pain remains unclear ^72^. The technique used in the present study has considerable potential in gaining a better understanding of the MOA, since the precise association between blood flow changes induced by ESCS and pain modulation is still under debate. Thus, the ability to accurately quantify blood flow in the spinal cord and/or brain networks both temporally and spatially and the increasing numbers of technologies available to modulate neural circuitry seem to be a logical clinical strategy to define MOA of pain and enhance therapeutic efficacy. This may require critically monitoring the spinal cord and/or brain during modulation with high spatiotemporal resolution and sensitivity systems. Additionally, fUSI could play a major role in identifying the MOA through which functional bidirectional connectivity across a functionally “complete” spinal injury can be reestablished with spinal neuromodulation techniques ^4^. Specifically, in the reorganization of neural networks that mediate spinal-supraspinal connectivity in children with cerebral palsy in response to spinal neuromodulation ^73^, and in the improvement of bladder function after paralysis ^74^. Overall, our study provides a high-resolution imaging technique to study the MOA of ESCS, and a tool to improve the efficacy of electrical stimulation by optimizing stimulation parameters based on the stimulation-evoked spinal cord hemodynamic signal.

### Evaluating the effectiveness of spinal cord stimulation protocols

One of the great advantages of fUSI is the ability to detect regional hemodynamic changes of only 2% without averaging over multiple trials. The ability to rely on the accuracy of a single-trial is necessary if one intends to use fUSI signal to monitor moment-to-moment hemodynamic changes. A single trial fUSI study was recently conducted to predict motor intention in NHPs that perform reaching and saccade (i.e., eye movements) to peripheral targets ^35^. By recording hemodynamic activity over the posterior parietal cortex (PPC), an association cortical region that encodes planning of actions, fUSI detected which effector the animals intended to use (eye or hand) and which direction they intended to move (left or right). These results show for the first time that fUSI is capable of imaging hemodynamics in large animals with enough sensitivity to decode the timing and goals of an intended movement. Building on this study, we designed a linear classifier that decodes fUSI signals of the spinal cord to accurately predict the effectiveness of a stimulation protocol within a single trial. Importantly, we showed that the best decoding accuracy of the spinal cord state is achieved using less than 30 s stimulation ON/OFF period data – i.e., around 24 s – to train the classifier. This finding suggests that hemodynamic changes induced by ESCS peak before turning off the stimulator and become less potent with time, impairing the decoding accuracy of the spinal cord state. This is also consistent with the observations from the ERA waveforms of the ΔSCBV that the pD signal in many regions returns to baseline activity before the stimulation is ceased (Fig. 4). Overall, these findings provide the first proof-of-concept that fUSI can detect changes of spinal cord hemodynamic responses induced by electrical stimulation within single-trials. It opens new avenues in neuromodulation for developing a real-time translational clinical spinal cord stimulation monitoring system.

### fUSI animal studies for revealing functional organization of spinal cord

While our study constitutes a novel and precise characterization of the hemodynamic signal of the human spinal cord, recent animal studies have utilized fUSI to monitor local hemodynamic responses to electrical or natural stimulation ^41–44^. The first spinal cord fUSI feasibility studies were performed in rat and swine animal models to explore the effects of stimulation-induced hemodynamic changes of the spinal cord ^41^. These studies show that fUSI offers better sensitivity in detecting and characterizing the spinal cord hemodynamic response during subthreshold-to-threshold motor response compared to electrophysiological assessment. A subsequent study evaluated the spinal cord hemodynamic responses to different stimulation parameters, such as frequency, electrode configuration and voltage intensity ^42^. By varying the stimulation parameters, the results provided strong evidence that distinct mechanisms are involved in spinal cord hemodynamic changes that correspond to different stimulation parameters. For instance, a strong association was found between spinal cord hemodynamics and EMG responses, but only at lower frequencies of electrical stimulation. Recently an animal study combined fUSI with electrical and natural stimulation of different categories of afferent fibers (Aβ, Aδ, and C fibers) to characterize, for the first time, the effects of peripheral nerve stimulation to the hemodynamics response of the rat spinal cord ^43^. The results show that the evoked hemodynamic responses are fiber-specific and located ipsilaterally in the dorsal horn of the rat spinal cord. Although these studies are insightful, they are based on animal models which limit the interpretability and the applicability of the findings to human populations. Our study is the first to successfully quantify fUSI activity in the human spinal cord with just a single trial of data. Compared to previous animal studies that used multi-trial detection of electrical spinal cord stimulation, detecting the effects of spinal cord stimulation from a single trial represents a new standard for closed-loop neuromodulation systems using fUSI technique.

## Conclusions

This study provides the first in-human quantitative evaluation of spinal cord hemodynamics using functional ultrasound imaging (fUSI). By combining fUSI and epidural electrical spinal cord stimulation (ESCS) it was possible to measure and characterize the spatially specific hemodynamics of the spinal cord. We describe a single-trial analysis to predict the effectiveness of a stimulation protocol to modulate the hemodynamic signal of the spinal cord. We also show that functional information useful for decoding the spinal cord state (i.e., stimulation ON vs. stimulation OFF) is primarily located in the small vessels within the imaging plane. As ongoing developments of fUSI hardware and software enable real-time imaging and processing, we can develop translational clinical monitoring systems that will help to evaluate the effectiveness of ESCS and other therapeutic neuromodulation strategies – a technology that currently does not exist. Overall, this work establishes fUSI as a promising platform for neuroscientific investigation with potential for profound clinical impact.

## Materials and Methods

### Patients and surgical procedure

Four patients underwent standard-of-care implantation of a SCS paddle lead (PentaTM model 3228) for treatment of chronic back pain conditions. Patients were recruited from the Department of Neurosurgery at the Keck School of Medicine of the University of Southern California (USC) and underwent a standard-of-care implantation of T10 partial laminectomy for insertion of a spinal cord stimulation paddle lead in the prone position, under general anesthesia. Once the location of the epidural lead was confirmed with fluoroscopy (lead spanning the T8-9 interspace), the fUSI probe was attached to an articulating arm and placed over the spinal cord for transverse imaging (Fig. 1A-C). The ESCS leads were then sterilely attached to an external pulse generator. Patients were eligible for inclusion if they were diagnosed with failed back syndrome. Informed consent was obtained from all patients after the nature of the study and possible risks were clearly explained. All experimental procedures were approved by the USC Institutional Review Board.

### Functional ultrasound imaging set-up

We utilized a fully featured commercial functional ultrasound acquisition system, *Iconeus One* (Iconeus, Paris, France) to acquire fUSI spinal cord images via a transducer probe inserted through laminar openings (Fig. 1C). A 128 elements linear array transducer (Fig. 1B, inner panel), with 15 MHz center frequency, 0.1 mm pitch (Vermon, Tours, France) was used to generate fUSI images with spatial resolution of 100 μm × 100 μm in-plane, slice thickness of 400 μm, field of view (FOV) of 12.8 mm (width) × 10 (depth) mm and maximum penetration depth of about 20 mm. The penetration depth is sufficient to image simultaneously almost the entire transverse cross-section of a human spinal cord, approximately 10-15 mm in diameter. The probe was fixed steadily throughout experiments with the field of view (FOV) transverse and intersecting the spinal cord central canal (Fig. 2A). To increase imaging signal-to-noise-ratio (SNR), we obtained each image from 200 compounded frames acquired at 500 Hz frame rate, using 11 tilted plane waves separated by 2° (i.e., from -10° to +10° increment by 2°). Images were acquired with a 5500 Hz pulse repetition frequency (PRF). Imaging sessions were performed using a real-time continuous acquisition of successive blocks of 400 ms (with 600 ms pause between pulses) of compounded plane wave imaging. The acoustic amplitudes and intensities of the fUSI sequence remained below the Food and Drug Administration limits for ultrasonic diagnostic imaging (FDA, 510k, Trace 3).

### Epidural electrical SCS parameters and stimulation protocol

A stimulation protocol was administered consisting of 10 ON-OFF cycles, with each cycle containing 30 s ON period (stimulation ON) and 30 s OFF period (stimulation OFF). The protocol involved bipolar stimulation at the middle contacts (spanning T8-T9 interspace), utilizing 40 Hz burst frequency, 300 μs pulse width and 6 mA amplitude.

#### Data analysis

##### Data pre-processing: Artifact removal and image stabilization

The Iconeus One acquisition system generates power Doppler (pD) intensity images pre-processed with built-in phase-correlation based sub-pixel motion registration ^75^ and singular-value-decomposition (SVD) based clutter filtering algorithms ^76^. These algorithms are used to separate tissue motion signal from blood signal to generate relative fUSI pD signal intensity images ^77^. To address potential physiological and motion artifacts unique to human spinal cord imaging, we adopt rigid motion correction techniques ^78^ that have successfully been used in fUSI ^35^ and other neuroimaging studies ^79–81^. This was combined with in-house developed breathing, high frequency smoothing filtering and image stabilization approaches, to ensure robustness and reliability of our data processing and analysis results.

### Statistical parametric maps and event-related average (ERA) waveforms

We leverage conventional statistical analysis in functional brain imaging to compute statistical parametric maps (SPM) to visualize regional spinal cord blood volume (SCBV) changes that are caused by ESCS. We identify regions with statistically significant changes (p < 0.05) on the fUSI signal between stimulation OFF and stimulation ON trials. Note that we used only the last 6 s of the stimulation OFF trials – i.e., 6 s prior to stimulation onset – to minimize the potential washout effect of the electrical stimulation. We selected 6 s because we found that it can take up to about 20 s for the hemodynamic signal to return to baseline activity after the stimulation was ceased (see Figs. 4B and 4C). The SPMs are derived based on the *Student’s t-test* (one side with false discovery rate [FDR] correction based on the number of voxels tested) for each pixel in the fUSI image. That is, each individual pixel derived its own p-value which indicates the level of significant ΔSCBV induced by ESCS. Finally, we generate event-related average (ERA) waveforms of the SCBV changes as a percentage change from baseline activity (i.e., average fUSI activity across 6 seconds preceding stimulation onset) for selected ROIs.

### Predict the effects of ESCS on spinal cord hemodynamic response at a single-trial level

We utilize conventional data analysis and interpretable machine learning (ML) techniques to examine whether evoked hemodynamic changes captured in single trial fUSI spinal cord images can be used to discriminate between stimulation ON and stimulation OFF states. We perform off-line decoding on the fUSI spinal cord images with a strategy that consists of three parts: 1) align the SCBV image time series with the stimulation status, 2) select features, reduce dimensionality, and discriminate classes, 3) cross validate and evaluate performance (Fig. 5). Each power Doppler (pD) image is flattened to create a 1D vector and is attached to a class label that represents the status of the stimulation (class 0: stimulation OFF and class 1: stimulation ON). A subset from each group is then separated into training (75%) and testing (25%) sets for cross-validation analysis. We use whole fUSI spinal cord images to perform the single-trial analysis. For dimensionality reduction and class separation, we employ classwise principal component analysis (cPCA) ^47^ and linear discriminant analysis (LDA) ^82^ respectively. We chose cPCA because of the small imaging sample size, large sparsity, and feature dimensions. It has successfully been used to reduce data sparsity and dimensionality while maintaining enough components to retain over 90% variance in imaging data such as ours ^47,83,84^. We integrate cPCA with linear discriminant analysis (LDA) to assess whether the cPCA-transformed fUSI images can predict an impending stimulation OFF (class 0) or stimulation ON (class 1). Mathematically the transformed feature for each trial can be represented by *f* = *T*_*LDA*_*Φ*_*CPCA*_(*d*), where *d* ∈ ℝ are the flattened images, *Φ*_*CPCA*_ is the piecewise linear cPCA transformation, and *T*_*LDA*_ is the LDA transformation. *Φ*_*CPCA*_ is physically related to space and thus can be viewed within the context of physiological meaning (Fig. 6D). We subsequently use Bayes rule to calculate the posterior probabilities of each class given the observed feature space. Because cPCA is a piecewise function, this is done twice, once for each class, resulting in four posterior likelihoods: *P*_*OFF*_(*OFF*|*f*^*\**^), *P*_*OFF*_(*ON*|*f*^*\**^), *P*_*ON*_(*OFF*|*f*^*\**^) and *P*_*ON*_(*ON*|*f*^*\**^) where *f*^*\**^ represents the observation, *OFF* and *ON* represent the posterior probabilities in the cPCA subspaces created with training data from stimulation OFF and stimulation ON trials, respectively. Finally, we store the optimal principal component (PC) vectors and the corresponding discriminant hyperplane from the subspace with the highest posterior probability. We then use these findings to predict the state of the spinal cord (i.e., stimulation OFF vs. stimulation ON) in the testing set. That is, we compute *f*^*\**^ from fUSI imaging data for each trial in the testing set to predict the state of the spinal cord. To cross-validate and test the robustness of the classifier models, we perform 100 iterations of bootstrapping randomized selection of 75% of the stimulation ON and OFF fUSI images for training our decoding model. The randomized bootstrap selection approach was utilized to accommodate the relatively small data size. All the data pre- and post-processing analysis were performed in MATLAB (Matworks, Natick, MA).

### Power Doppler (pD) quantiles

To evaluate the source of hemodynamic information content relevant for classifying ESCS ON and OFF states, we ranked ordered voxels by their mean pD intensity, which is derived from averaging 6 s of fUSI recordings prior to the first stimulation trial and segmented by deciles resulting in a spatial map of ranked deciles (Fig. 7A). Higher deciles correspond to higher pD intensity and hence higher blood flow in the spinal cord. The voxels within each decile were unique and did not overlap with other deciles. Using voxels from each segmented decile, we performed cross-validated classification analysis, as described before, to evaluate the classification accuracy across deciles.

## Data availability

The datasets generated and analyzed during the current study are available from the corresponding authors on reasonable request.

## Acknowledgments

We thank the participants that made this study possible. We acknowledge J. Gibbons and S. Oviedo for administrative support. This work was supported by “The Center of Neurorestoration” at the University of Southern California and the “Marlan and Rosemary Bourns College of Engineering” at the University of California Riverside through start-up funding.

## Author Contributions

D.J.L., J.R., V.R.E., C.L. and V.N.C conceived the study.; D.J.L. performed the surgeries; D.J.L., W.C. and A.A. acquired the functional ultrasound data; K.A.A. and V.N.C. performed the functional ultrasound data processing, statistical and machine learning decoding analysis; K.A.A. and D.J.L. drafted the manuscript with substantial contribution from J.R., E.I.K, V.R.E., C.L. and V.N.C; All authors edited and approved the final version of the manuscript; V.R.E, C.L. and V.N.C. supervised the research.

---

**Fig. S2.**
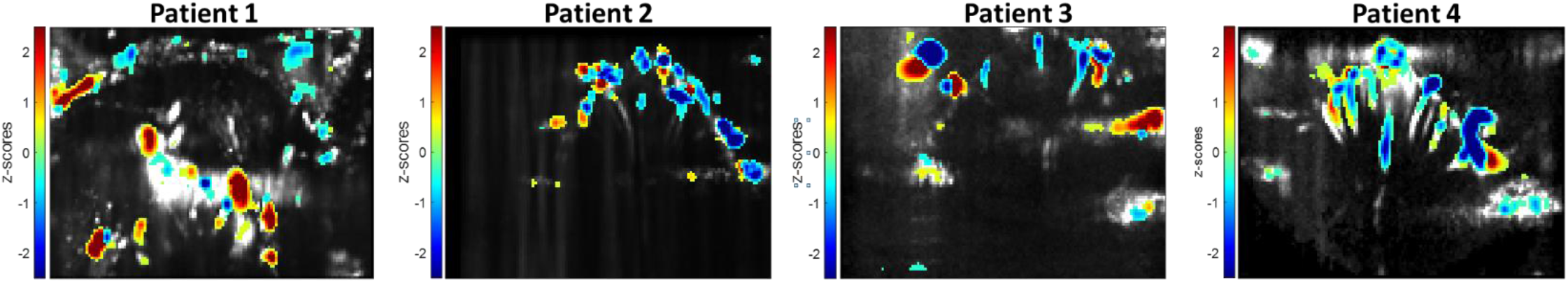
Statistical parametric maps (SPM) without correcting for washout effects from the electrical stimulation. Statistical map shows localized areas with significantly higher SCBV with respect to baseline (one-sided t test of area under the curve, p < 0.05, false discovery rate [FDR] corrected for number of pixels in image). We do not correct for potential washout effect from the electrical stimulation – i.e., all 30 s of acquisition were used from the stimulation OFF trials.

